# Genetic evidence for signal transduction within the *Bacillus subtilis* GerA germinant receptor

**DOI:** 10.1101/2021.10.28.466341

**Authors:** Jeremy D. Amon, Lior Artzi, David Z. Rudner

## Abstract

Bacterial spores can rapidly exit dormancy through the process of germination. This process begins with the activation of nutrient receptors embedded in the spore membrane. The prototypical germinant receptor in *Bacillus subtilis* responds to L-alanine and is thought to be a complex of proteins encoded by the genes in the *gerA* operon: *gerAA*, *gerAB*, and *gerAC*. The GerAB subunit has recently been shown to function as the nutrient sensor, but beyond contributing to complex stability, no additional functions have been attributed to the other two subunits. Here, we investigate the role of GerAA. We resurrect a previously characterized allele of *gerA* (termed *gerA\**) that carries a mutation in *gerAA* and show it constitutively activates germination even in the presence of a wild-type copy of *gerA*. Using an enrichment strategy to screen for suppressors of *gerA\**, we identified mutations in all three *gerA* genes that restore a functional receptor. Characterization of two distinct *gerAB* suppressors revealed that one (*gerAB-E105K)* reduces the GerA complex’s ability to respond to L-alanine, while another (*gerAB-F259S*) disrupts the germinant signal downstream of L-alanine recognition. These data argue against models in which GerAA is directly or indirectly involved in germinant sensing. Rather, our data suggest that GerAA is responsible for transducing the nutrient signal sensed by GerAB. While the steps downstream of *gerAA* have yet to be uncovered, these results validate the use of a dominant-negative genetic approach in elucidating the *gerA* signal transduction pathway.

**Importance:** Endospore formers are a broad group of bacteria that can enter dormancy upon starvation and exit dormancy upon sensing the return of nutrients. How dormant spores sense and respond to these nutrients is poorly understood. Here, we identify a key step in the signal transduction pathway that is activated after spores detect the amino acid L-alanine. We present a model that provides a more complete picture of this process that is critical for allowing dormant spores to germinate and resume growth.

## Introduction

In response to nutrient deprivation, many bacterial species from the orders Bacillales and Clostridiales differentiate into metabolically dormant and highly resilient endospores (hereafter referred to as “spores”) (1, 2). Spore-forming organisms are ubiquitous in the environment and include many medically, agriculturally, and technologically important species. These include the causative agents of anthrax and tetanus, the pesticide-producing *Bacillus thurengiensis*, and *Bacillus cereus*, an agent of food-borne illness (3, 4). Spores have the remarkable ability to resist biological, chemical, and physical assaults, and can remain dormant for decades(3, 5). And yet despite this ability to remain inert under extreme conditions, they can rapidly exit dormancy upon exposure to specific nutrients through a process called germination (6, 7).

Germination relies on a finely tuned mechanism of environmental sensing and subsequent response. The process is initiated by germinant receptors embedded in the spore inner membrane, which are triggered by a range of small molecules including amino acids, nucleosides, and sugars (8). In *Bacillus subtilis*, the prototypical germinant receptor is encoded by the genes of the *gerA* operon – *gerAA*, *gerAB*, and *gerAC* – and can induce germination in response to L-alanine (9–11). Recent work has established that nutrient sensing involves a binding pocket in the core of the polytopic membrane protein GerAB (Artzi, *et al*., manuscript under review). The roles of GerAA and GerAC in the germination pathway remain unknown. The next steps have been described physiologically, but mechanistic details remain lacking. First, monovalent ions and stores of the small molecule dipicolinic acid (DPA) in complex with Ca^2+^ are released from the spore core, allowing partial rehydration (12, 13). Next, two functionally redundant cell wall hydrolases, SleB and CwlJ, are activated and digest the specialized spore peptidoglycan, known as the cortex (14, 15). CwlJ is thought to be activated by DPA, but the precise mechanism of activation remains unclear, as does the mechanism of activation of SleB (16, 17). Degradation of the spore cortex by SleB and CwlJ allows complete rehydration of the core, resumption of metabolic processes, and outgrowth of the vegetative cell.

Mis-regulation of this process can be catastrophic for the organism. Indeed, any spore that is unable to germinate is essentially dead (18). Conversely, spores that germinate too readily risk emerging into a compromised environment (19). These two extremes highlight the twin therapeutic potentials of studies on germination: spores that can be prevented from germinating pose no threat, and those that can be forced to germinate can be easily treated with antibiotics. But much remains to be understood about the germination pathway and its potential Achilles’ heels before it can be exploited for medical or industrial purposes. With the L-alanine binding pocket in GerAB now identified, the question becomes what happens next? How is the signal transduced? And what, if any, are the roles of GerAA and GerAC in information transduction?

Due to the challenges of reconstituting the GerA membrane complex in vitro and establishing an assay for signal transduction, these questions have gone largely unanswered. However, the presence of a receptor-ligand interaction at the head of a relatively unexplored signaling pathway is an alluring prospect for classical genetic analysis. Theoretically, constitutively active mutants of the *gerA* receptor that prematurely trigger premature spore germination can be exploited. Suppressors of such constitutively active mutants can be selected because they will regain their heat resistance through inhibition of *gerA*-induced germination. These suppressors have the potential to reveal steps downstream of nutrient detection.

Previously published work uncovered a potential candidate for such analysis. An allele of *gerAA*, *gerAA-113*, was found to have a single amino acid change of a proline at position 326 to serine. This mutation resulted in the production of phase-dark spores at the end of sporulation. The authors hypothesized that this phenotype was a result of the receptor being triggered spontaneously (20). Here, we resurrect *gerAA-113* and show that it is a dominant-negative hyperactive allele and thus well suited to genetic analysis. Using this allele, we identified suppressors that map to the *gerA* locus. The characterization of two *gerAB* suppressors provide evidence that GerAA is responsible for transducing the nutrient signal sensed by GerAB. We discuss these alleles in the context of an AlphaFold2-predicted structure (21, 22) of the GerA complex.

## Results

### *gerAA(P326S)* triggers DPA release and SleB activation

We began by characterizing the *gerAA(P326S)* allele. We reasoned that if this allele is hypermorphic, it should be dominant-negative and result in phase dark spores in a merodiploid strain carrying the native copy of *gerA*. We rebuilt the *P326S* mutation into the *gerA* operon and inserted it at an ectopic locus (*ycgO*) in the genome. For the purposes of this manuscript, this allele, *ycgO::gerAA(P326S)-gerAB-gerAC,* is referred to as *gerA\**. As can be seen in Figure 1A, sporulating cells harboring *gerA\** and the native *gerA* locus produced an abundance of phase-dark spores. Furthermore, when sporulated cultures of the *gerA*/gerA^+^* merodiploid were subjected to heat treatment to kill vegetative cells and heat-sensitive spores, there was a corresponding drop in spore viability to approximately 3% of wild-type. Importantly, these phenotypes were not observed in a strain containing an additional copy of the wild-type *gerA* operon at the same ectopic locus. We conclude that *gerA\** is a hyperactive dominant-negative allele.

**Figure 1:**
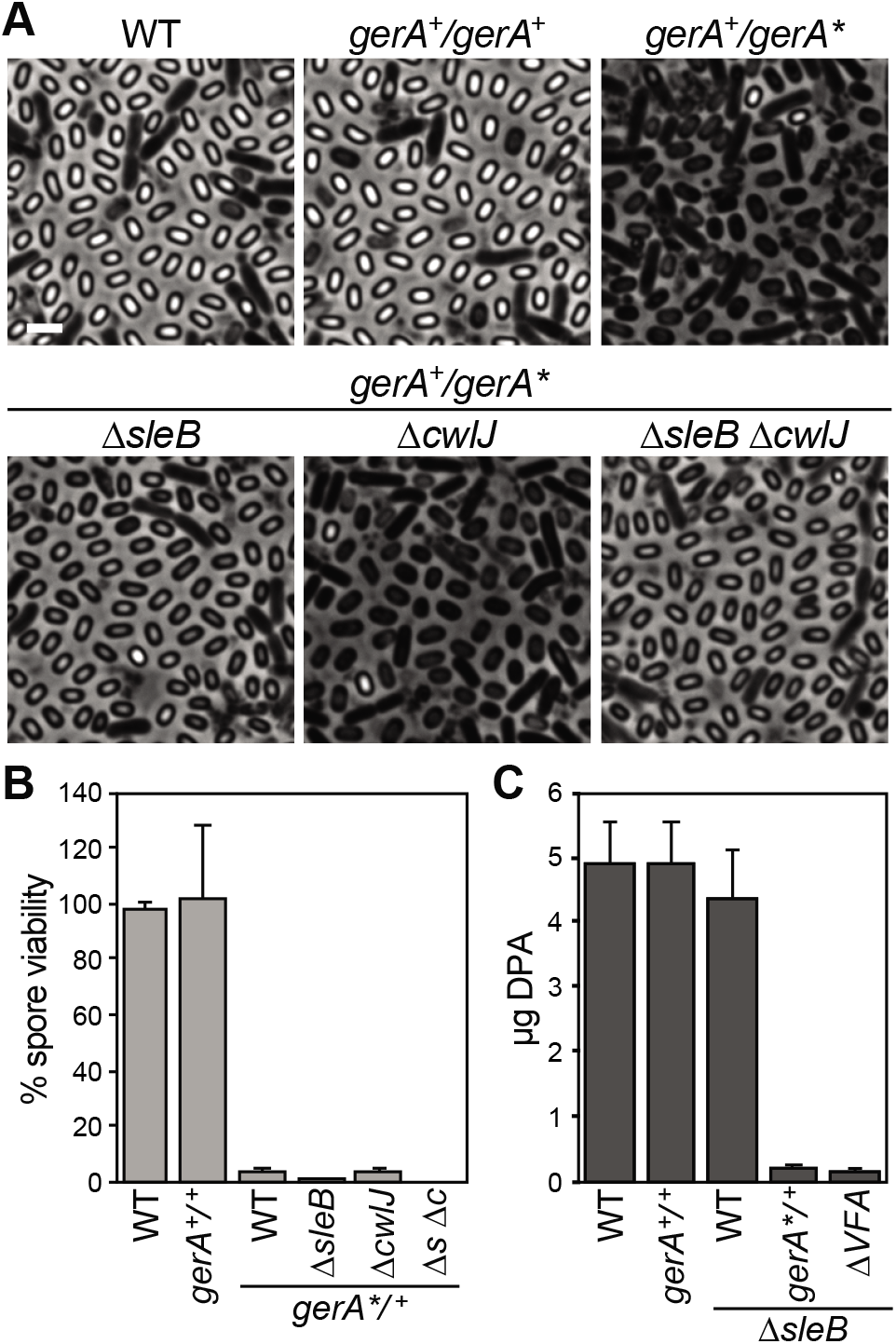
GerA* triggers DPA release and SleB activation. (A) Phase-contrast micrographs of the indicated strains sporulated by nutrient exhaustion for 30 h. *gerA^+^/gerA^+^* is a merodiploid strain with two copies of the wild type *gerA* locus. *gerA^+^/gerA\** is a merodiploid strain with a wild type *gerA* locus and a mutant *gerA\** operon at an ectopic genomic locus. Experiments were performed in biological triplicate; representative images are shown. Scale bar is 2 μm. (B) Sporulated cultures in (A) were heat treated (80°C for 20 min), and serial dilutions were plated on LB to assess heat-resistant colony forming units. Wild type spore viability (3.3×108 CFU/ml) was set to 100%. *Δs Δc* − *ΔsleB ΔcwlJ*. Error bars indicate standard deviation, n=3. (C) Phase-grey and -bright spores were purified from sporulated cultures in (A). Spores were boiled to release DPA. DPA was then quantified using TbCl_3_ compared to standards. Values are reported as micrograms of DPA released from 1ml of purified spores adjusted to OD600=1. *ΔVFA* − *ΔspoVFA*. Error bars indicate standard deviation, n=4.

Previous studies have shown that premature germination is dependent on the cell wall hydrolase SleB (23). To confirm that *gerA\** was inducing premature germination, we analyzed the *gerA*/gerA^+^* merodiploid in a Δ*sleB* background. As anticipated, this strain produced phase-gray spores instead of phase-dark spores. Furthermore, the absence of SleB in the *gerA*/gerA^+^* strain caused a further drop in spore viability to approximately 0.2% of wild-type. By contrast, the absence of the functionally redundant cell wall hydrolase CwlJ did not alter the phase-dark phenotype nor did it alter spore viability. Thus, *gerA*\* cause premature germination through the activation of SleB.

The appearance of phase-gray spores is thought to reflect the failure to accumulate and retain DPA and the partial hydration of the core. To investigate whether *gerA*\* spores accumulate and retain DPA, we purified spores from the *gerA*/gerA^+^ ΔsleB* mutant and quantified the intracellular stores of DPA. As can be seen in Figure 1C, wild-type, *gerA^+^/gerA^+^,* and Δ*sleB* spores contained 4-5 μg of DPA per 1 OD_600_ equivalent. By contrast, *gerA^+^/gerA* ΔsleB* spores contained 0.24 μg. This low level was comparable to the 0.16 μg present in spores from *ΔspoVFA ΔsleB* spores, which lack the DPA synthesis machinery. Altogether, these data support the idea that sporulating cells that produce GerA receptors with GerAA(P326S) trigger SleB activation and the release of DPA during spore formation.

### An enrichment screen for suppressors of *gerA\** identifies mutations throughout the gerA operon

To probe the germination pathway, we sought to identify mutations that suppress the premature germination caused by *gerA*\*. Due to the presence of other functional germinant receptors, the most common suppressors of *gerA\** were expected to be mutations that inactivate *gerAA*(P326S). Accordingly, to select for informative suppressors of *gerA*,* we deleted the native *gerA* locus and the operons encoding the other four Ger receptors (*gerB, gerK*, *yfk, ynd*). This strain, referred to as *Δ5*, is unable to respond to germinants, with a resultant spore viability of 0.24% of wild-type (Figure 2A). Importantly, an ectopic copy of the wild-type *gerA* locus can complement the *Δ5* strain, restoring spore viability to 82%. By contrast, spore viability of the *Δ5* strain harboring *gerA\** was 3.5%. As anticipated, *gerA\** continued to inappropriately cause DPA release and SleB activation when it was the sole germinant receptor (Supplemental Figure 1). The *Δ5* strain therefore creates a restrictive genetic background. Suppressors of *Δ5 gerA\** cannot simply inactivate *gerA\** or else they would be unable to germinate. Rather, suppressors must maintain function while abrogating premature activation.

**Figure 2:**
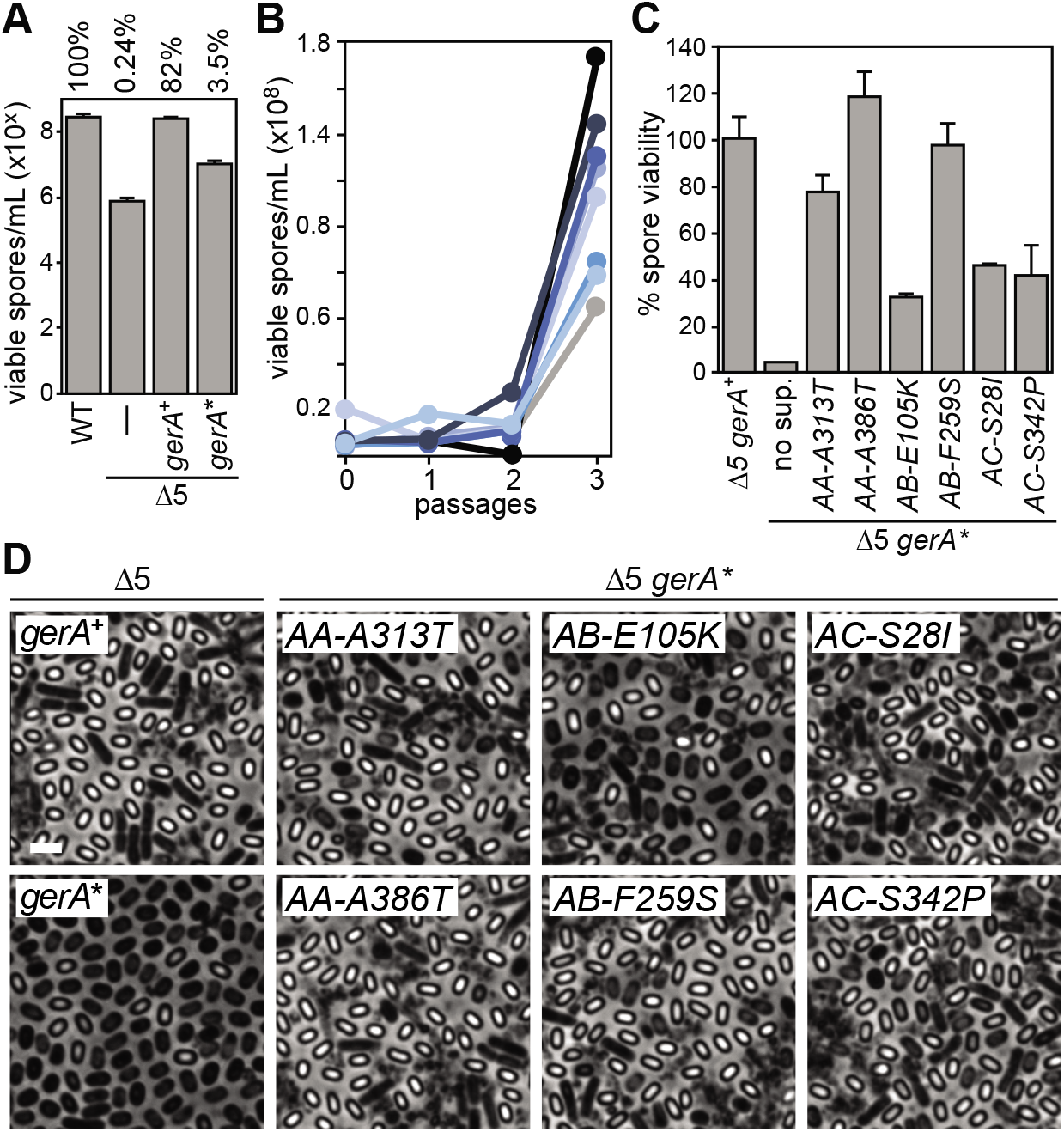
An enrichment screen for suppressors of *gerA\**. (A) Cultures were sporulated by nutrient exhaustion and heat-treated (80°C for 20 min). Serial dilutions were plated on LB to assess heat-resistant colony forming units. Wild type spore viability (3.0 x108 CFU/ml) was set to 100%. *Δ5* – *ΔgerA ΔgerBB ΔgerKB ΔyndE ΔyfkT*. Values are reported on a logarithmic scale for clarity. Error bars indicate standard deviation, n=5. (B) Cultures of *Δ5 gerA\** cells were sporulated by nutrient exhaustion and heat-treated. A sample was taken to assess heat-resistant colony forming units by plating serial dilutions on LB. Another sample of ~10^6^ viable spores was then used to inoculate fresh cultures, which allowed the spores to germinate, outgrow, and re-sporulate upon nutrient exhaustion. This process was repeated until spore viability increased. A sample of eight independent lineages are shown. Values are reported on a linear scale for clarity. (C) Cultures were sporulated overnight and heat treated. Serial dilutions were plated on LB to assess heat-resistant colony forming units. *Δ5 gerA^+^* spore viability (2.2 x108 CFU/ml) was set to 100%. Error bars indicate standard deviation, n=3. (D) Micrographs of sporulated cultures from (C). Representative images from three biological replicates are shown. Scale bar is 2 μm.

To select for suppressors of *Δ5 gerA\**, we used a serial enrichment strategy. Overnight sporulation cultures were heat treated to kill vegetative cells and prematurely germinated spores. A sample of the culture was taken to monitor spore viability by plating for CFUs. Approximately 10^6^ viable, heat-resistant spores were then used to inoculate fresh sporulation medium. This process was repeated until suppressors overtook the culture and resulted in an increase in viable spores, which almost always occurred after the third passage (Figure 2B). Cultures were then streak-purified and individual colonies were screened for the suppressive phenotype.

In total, 83 suppressors of *Δ5 gerA\** were identified and further characterized. We began by backcrossing the *gerA*\* locus into the Δ5 mutant to determine whether the suppressors were linked to *gerA*\*. Strikingly, all 83 suppressors mapped to the *gerA*\* locus. In total, 46 unique mutations were identified in 38 codons spanning all three genes in the operon (Supplementary Table 1). Figure 2C shows spore viability of six representative suppressors within *gerAA, gerAB,* and *gerAC*, with suppression of the *gerA\** allele ranging from a 10- to 30-fold increase in spore viability. Phase-contrast microscopy of overnight sporulating cultures confirmed suppression of premature germination, with the proportion of phase-bright spores increasing as suppression improved (Figure 2D).

### *gerAB* mutants attenuate the germination signal upstream of *gerA\**

Recent studies have established an L-alanine binding pocket in the core of the GerAB protein (Artzi, *et al*., manuscript under review). The existence of *gerA\** suppressors in *gerAB* presented a possibility to investigate how the GerAA subunit is linked to the binding pocket. In one model, *gerAA* plays a purely structural role and stabilizes the complex. In another, *gerAA* acts “upstream” of L-alanine binding, either by controlling access to the binding pocket or shaping the binding pocket to tune specificity or affinity. In a third model, *gerAA* acts “downstream” of L-alanine binding, transducing and/or processing the signal and passing this information to other parts of the complex or to other proteins.

To explore these models, we rebuilt our *gerAB* suppressor mutations and introduced them at an ectopic genomic locus in a strain that lacked *gerAB* and contained inactivating alleles of *gerB, gerK, yfk,* and *ynd (Δ4 ΔgerAB*). As seen in Figure 3A, a wild type copy of *gerAB* inserted at the ectopic locus complemented the Δ*4 ΔgerAB* strain. The absence of a complementing allele phenocopied the Δ*5* strain, with spore viability at 0.2% of wild-type. Six *gerAB* suppressors (*R107Q, R107W, W253L, G266D, G266S, I267R*) inserted at the same ectopic locus partially complemented the Δ*4 ΔgerAB* strain, with spore viability restored to 15-40% of wild-type. One explanation for why hypomorphic *gerAB* mutants suppress *gerAA(P326S)* is that the hyperactive GerA* receptor complex can respond to low levels of L-alanine present during sporulation. This concentration is not sufficient to induce germination of wild-type GerA but can further stimulate germination of GerA*. In this context, hypomorphic *gerAB* alleles suppress *gerA*\* by dampening this response.

**Figure 3:**
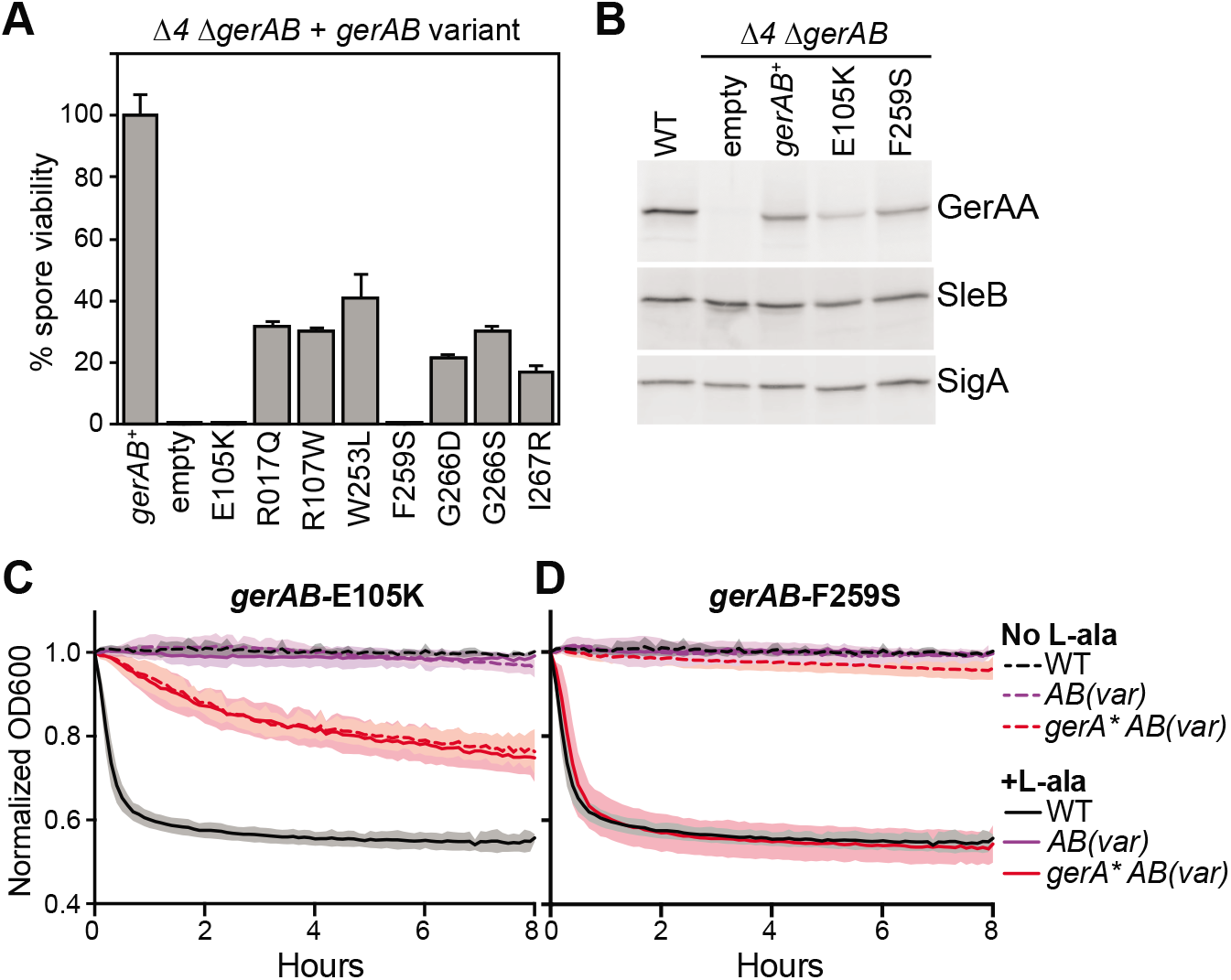
Analysis of GerAB suppressor mutants. (A) The indicated strains were sporulated overnight and heat treated. Serial dilutions were plated on LB to assess heat-resistant colony forming units. *Δ4* – *ΔgerBB ΔgerKB ΔyndE ΔyfkT*. *gerAB+* spore viability (3.5 x108 CFU/ml) was set to 100%. Error bars indicate standard deviation, n=3. (B) Immunoblots of *gerAB* variants. Cultures were sporulated overnight. Phase-grey and phase-bright spores were purified using lysozyme and SDS. Spores were physically disrupted, and lysates were subjected to SDS-PAGE followed by immunoblot analysis to detect the presence of GerAA. SleB and SigA were analyzed to control for loading. Representative immunoblots are shown, n=3. (C-D) Germination of *gerAB* variants. Cultures were sporulated overnight, and phase-bright spores were purified using density gradients. Spores were resuspended in 96-well plates in the presence or absence of L-alanine and agitated for 8 hours at 37°C. Optical density at time zero was normalized to 1, and subsequent measurements were taken every two minutes. Error bars indicate standard deviation of three biological replicates. *AB(var)* – indicates *gerAB(E105K)* or *gerAB(F259S)* in (C) or (D), respectively.

In addition to the hypomorphic alleles, we identified two *gerAB* suppressors (*E105K* and *F259S*) that failed to complement Δ*4 ΔgerAB* mutant, with spore viability of 0.2%. This loss-of-function phenotype was not due to destabilization of the germination complex, as the levels of GerAA in spores derived from both mutants were similar to wild-type GerAB (Figure 3B). To directly assess germination in these GerAB mutants, we purified phase-bright spores and monitored germination kinetics after addition of L-alanine. As previously shown, wild-type spores are stable in the absence of nutrients but germinate within 30 minutes after the addition of 1 mM L-alanine, as assayed by a drop in optical density (24). As anticipated, Δ*4 ΔgerAB* strains harboring either *gerAB(E105K)* or *gerAB(F259S)* were stable over 8 hours and were unresponsive to the addition of L-alanine (Figures 3A-B). Collectively, these data indicate that *gerAB(E105K)* and *gerAB(F259S)* are stable, null alleles of *gerAB*.

The fact that these *gerAB* null alleles were able to form viable spores when combined with *gerAA(P326S)* led us to hypothesize that *gerAA* acts downstream of *gerAB*. To characterize this genetic interaction further, we purified phase-bright spores from the *gerAB* mutants in the *gerA\** background. Interestingly, the *gerAB(E105K) gerA\** spores germinated slowly but steadily over 8 hours in the absence of L-alanine (Figure 3C). Microscopic examination of the purified spores over this 8-hour period confirmed this spontaneous germination and transition to phase-dark spores without the addition of L-alanine (Supplemental figure 2). Importantly, the addition of L-alanine did not change the germination rate of these spores. These data indicate that *gerAB(E105K)* is unresponsive to L-alanine, and that *gerAA(P326S)* bypasses this defect by germinating in a constitutive manner.

Similarly, purified spores carrying *gerAA(P326S)* in combination with *gerAB(F259S)* germinated in the absence of L-alanine, albeit very slowly (Figure 3D). Phase-contrast microscopy confirmed this subtle phenotype, revealing a subset of spores that had spontaneously germinated in the absence of L-alanine (Supplemental figure 2). However, in contrast to *gerAB(E105K)*, spores carrying *gerAA(P326S)* and *gerAB(F259S)* are responsive to L-alanine, germinating as rapidly and fully as WT. We conclude that GerAB(F259S) is responsive to L-alanine but somehow dampens the signal to the point of loss-of-function. However, in the presence of GerAA(P326S), the dampened signal is restored leading to a wild-type germination response.

### *gerAB(F259S)* blocks germination signal emanating from the GerAB binding pocket

To further explore whether the F259S substitution in GerAB dampens the nutrient signal emanating from the binding pocket, we turned to other hyperactive *gerA* alleles. A recent study (Artzi, *et al*., manuscript under review) found that mutating key binding pocket residues in *gerAB* can mimic L-alanine binding and result in premature germination. One such allele, *gerAB(T287L)*, produces phase-dark spores and results in 21% spore viability. To test whether the F259S substitution could dampen the premature germination signal resulting from *gerAB(T287L)*, we combined the two mutations in *gerAB* and tested spore viability. We found that F259S prevented premature germination triggered by the T287L substitution, producing phase bright spores (Figure 4). Furthermore, consistent with *F259S* being downstream of *T287L*, we found that spore viability of the *gerAB(F259S, T287L)* double-mutant phenocopied the *gerAB(F259S)* substitution, with 0.17% spore viability. This contrasts with the interaction between *gerAA(P326S)* and *gerAB(F259S)*, which results in 97% spore viability (Figure 2C). Similar results were found when analyzing another allele that activates germination by mimicking L-alanine binding, *gerAB(V101F)* (Supplemental Figure 3). Taken together, we conclude the *gerAB(F259S)* mutant uncovers a step downstream of nutrient sensing in the spore germination pathway.

**Figure 4:**
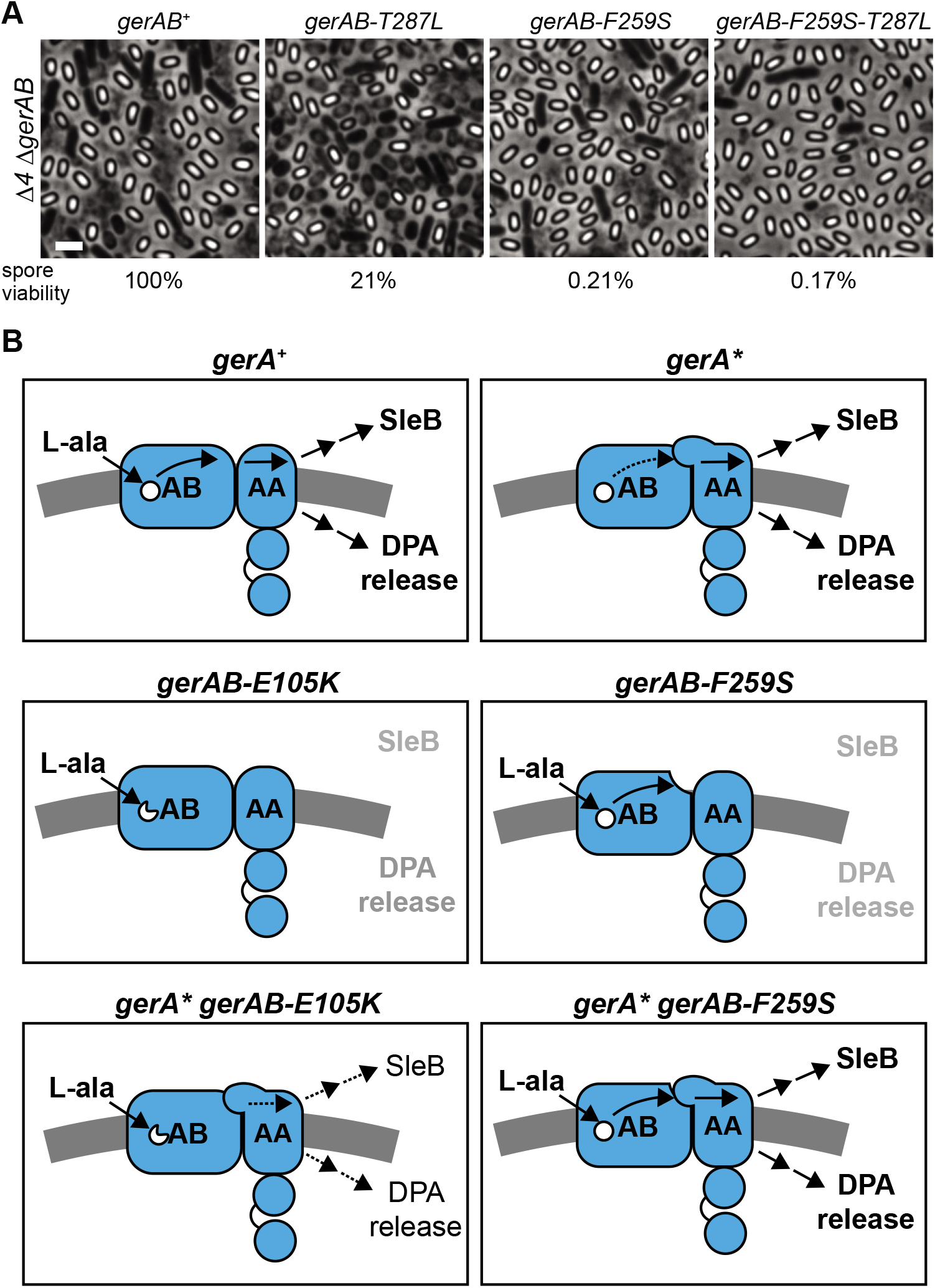
Epistatic analysis of *gerAB-F259S* and *gerAB-T287L*. (A) Cultures were sporulated by nutrient exhaustion. Representative phase-contrast micrographs of three biological replicates are shown. Scale bar is 2μm. Cultures were heat-treated (80°C for 20 min) then serially diluted and plated on LB to assess heat-resistant colony forming units. *gerAB+* spore viability (3.7 x108 CFU/ml) was set to 100%. (B) Model of signal transduction within the GerA complex. GerAC has been omitted for clarity.

### AlphaFold-predicted structure of the GerA complex reveals a putative signal transduction domain

The *gerAA(P326S)* and *gerAB(F259S)* variants are mutually suppressive and together restore the wild-type L-alanine response. This led us to consider the possibility that the two mutations alter a contact point between GerAA and GerAB. To explore physical interactions between GerAA and GerAB, we assembled a predicted structure of the GerA complex using AlphaFold2.0 (21, 22). The AlphaFold algorithm predicts that GerA forms a complex (Supplementary Figure 4A). GerAA and GerAB are predicted to lie adjacent to one another, with transmembrane helix four of GerAB predicted to be in direct contact with the sixth transmembrane helix of GerAA (Figure 5A). Furthermore, GerAC is predicted to contact both GerAA and GerAB on their extracellular surfaces.

**Figure 5:**
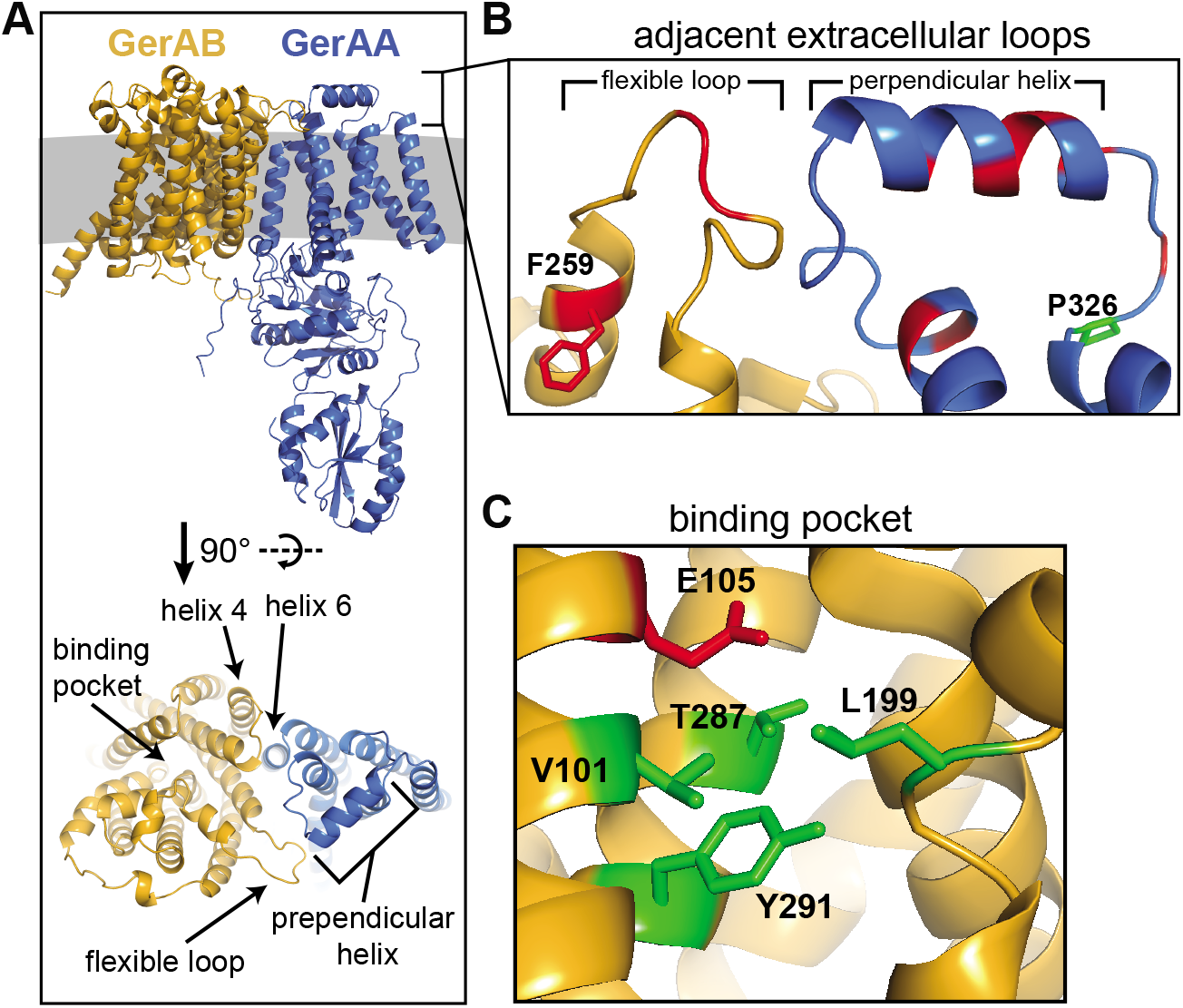
Predicted structures of GerAA and GerAB. (A) Top – Alphafold2-predicted structures of GerAA and GerAB in complex, situated in the inner spore membrane (grey). Bottom – Predicted structures rotated 90 degrees for a top-down view. Helix 4 of GerAB and helix 6 of GerAA are predicted to form a contact interface between the two proteins. The binding pocket of GerAB is visible, as is the proximity of the flexible loop in GerAB and the perpendicular helix in GerAA. (B) View of the predicted interface between the flexible loop of GerAB and the perpendicular helix of GerAA. Residue F259 of GerAB is shown in red and residue P326 is shown in green. Other residues that, when mutated, were able to suppress *gerAA(P326S)* are highlighted in red. (C) View of the L-alanine binding pocket in GerAB. Residue E105K is shown in red. Residues that have been previously shown to constitute the binding pocket (V101, T287, L199, Y291) are shown in green.

In addition to these contacts, the algorithm predicted that the first extracellular loop of GerAA and the fourth extracellular loop of GerAB are in proximity. On GerAA, this extracellular loop consists of a short helix oriented perpendicularly to the transmembrane helices (Figure 5B). Residue P326 of GerAA is predicted to be an anchor point of this loop. Interestingly, 13 of the 31 unique hits that mapped to *gerAA* were found on this extracellular loop or its anchor points, suggesting that stabilizing this loop may be critical to restoring GerAA function (Supplementary Table 1). On GerAB, extracellular loop four is a region of low confidence, as measured by predicted local distance difference test (plDDT), indicating that this loop may be flexible or unstructured (Supplemental Figure 4C). Residue F259 of GerAB is predicted to be at an anchor point of this loop. Five of the ten unique mutants that mapped to *gerAB* were found on this flexible loop or its anchor points, indicating that this loop may also be critical to restoring GerAA function (Figure 5B). These findings raise the possibility that an interaction between these extracellular loops could serve to transduce the germination signal emanating from the binding pocket in GerAB.

Lastly, to better understand how *gerAB(E105K)* was affecting nutrient detection, we analyzed the L-alanine binding pocket of GerAB. Residue E105 is predicted to be in close proximity to the four residues that have been previously shown to constitute the GerAB binding pocket (V101, L199, T287, Y291). The predicted structure suggests that GerAB(E105K) is unresponsive to L-alanine either because it distorts the binding pocket or prevents it from transmitting signal to the rest of the complex.

## Discussion

We have shown that *gerAA(P326S)* inappropriately activates germination by triggering both DPA release and SleB activation. We selected for suppressors of this allele and found 46 unique mutations in the *gerA* operon. Of these mutations, ten mapped to *gerAB*, two of which were of particular interest. The first, *gerAB(E105K)*, failed to germinate in response to L-alanine. In combination with *gerAA(P326S)*, *gerAB(E105K)* remained unresponsive to L-alanine and germinated constitutively at a slower rate than *gerAA(P326S)* itself. Position E105 is predicted to be near the binding pocket in the core of GerAB. The second suppressor of interest, *gerAB(F259S)*, also failed to germinate in response to L-alanine and furthermore could inhibit premature germination induced by mutations in the nutrient binding pocket. In combination with *gerAA(P326S)*, *gerAB(F259S)* regained its responsiveness to L-alanine, and germinated constitutively, albeit at a very slow rate. Position F259 is predicted to be at the base of a flexible, extracellular loop of GerAB.

Taken together, these results inform a model of signal transduction within the GerA receptor complex (Figure 4B). In the wild-type case, the complex is in the “off” state yet poised to respond. When L-alanine binds the pocket in the core of GerAB, a signal is transduced to the extracellular junction between GerAB and GerAA. GerAA then somehow transduces this signal to activate SleB and release DPA. When *gerAA(P326S)* is present, the extracellular junction between GerAB and GerAA is hypersensitive to signal from the binding pocket, and low levels of L-alanine in the sporangia are enough to push the complex into the “on” state. Inhibiting nutrient recognition, as with *gerAB(E105K)*, reveals the baseline constitutive activity of the *gerAA(P326S)* allele. In contrast, dampening the hyperactive extracellular junction, as with *gerAB(F259S)*, reverts the region to near-wild type, allowing the GerA complex to largely remain off yet responsive to L-alanine. When *gerAB(E105K)* or *gerAB(F259S)* are present individually, they shut down GerA completely – the first by inactivating the binding pocket, the second by dampening the extracellular junction to the point of preventing signal transduction.

Previous studies have presented evidence that GerA is likely poised on a knife’s edge to respond to L-alanine (23, 25). This allows GerA to respond quickly to germinant but results in a sizable fraction of wild-type spores that will prematurely germinate due to low levels of L-alanine in the sporangia. Our analysis of *gerAA(P326S)* supports this model and offers two mechanistic explanations for how GerA has tuned its sensitivity. One, as previously shown, is by the more direct method of altering the binding pocket. The other is by changing the nature of the interaction between the extracellular loops of GerAA and GerAB. Interestingly, the studies mentioned above found that other germinant receptors, GerB and GerK, played no role in premature germination. This could be due to lower levels of GerB- and GerK- specific germinants in the sporangia or due to different sensitivities of the receptors. Analyses of the interacting extracellular region in other germinant receptors could elucidate how spores set their sensitivities to changes in their environments.

The mechanism by which *gerAA* activates downstream events remains unknown. However, the presence of a cluster of mutants predicted to reside on the extracellular region of GerAA most distal to GerAB raises the possibility that the germination signal culminates in an event at this location (GerAA residues V369, E370, A371). One possibility is that this position may be an interaction domain with another factor. Alternatively, this region could be the site of a GerAA-dependent alteration of some physical property of the spore. In the context of this model, either would in turn trigger DPA release and SleB hydrolase activation.

Selecting against the dual activities of the GerA complex – DPA release and SleB activation – resulted in mutants that suppress both functions simultaneously. The fact that all 83 suppressors of *gerA\** were in the *gerA* operon suggests that there are no protein factors downstream of GerA capable of triggering both activities. Given these 83 suppressors, if we consider mutations within the *gerA* operon and external to the *gerA* operon as binary outcomes, the genetics argue with >98% confidence that finding a dual-function protein factor downstream of GerA has less than 1 in 20 odds. We therefore favor a model in which the GerA complex itself, and not some external protein factor, is the branch point between DPA release and SleB activation. Still, the presence of such a factor remains possible. Our screen has validated the use of a dominant-negative genetic approach to investigating GerA. Future studies will use similar screens to find such factors and saturate the mutational landscape of GerA.

## Materials and Methods

### General Methods

All *B. subtilis* strains were derived from the auxotrophic (*trpC2*) strain 168 (26). To obtain spores, sporulation was induced by nutrient exhaustion. Cells were grown in liquid Difco Sporulation Medium (DSM) at 37 °C with agitation for 24-30 hours. Alternatively, cells were grown on solid DSM agar at 37 °C for 96 hours. Spore viability was determined by comparing the total number of heat-resistant (80 °C for 20 min) colony forming units (CFUs) as a percentage of wild-type heat-resistant CFUs. Deletion mutants were derived from the *Bacillus* knock-out (BKE) collection (27) or were generated by isothermal assembly of PCR products (28) followed by direct transformation into *B. subtilis*. All BKE mutants were back-crossed twice into *B. subtilis* 168 before assaying and prior to antibiotic cassette removal. Antibiotic cassette removal was performed using a temperature-sensitive plasmid that constitutively expresses Cre recombinase (29). All strains were constructed using a 1-step competence method. Tables of strains, plasmids, and primers used in this study can be found in supplemental information.

### Genetic screen details

Fresh colonies of BJA177a (Δ*5 gerA\**) were suspended in 3ml DSM and incubated at 37 °C for 24 hours with agitation. Cultures were heat treated at 80 °C for 20 min. 100µl were removed for dilutions and CFU plating. An additional 100µl (~10^6^ viable spores) were removed and used to inoculate fresh 3ml DSM cultures. This process was repeated until spore viability had appreciably increased, indicating that suppressors had overtaken the cultures. These cultures were genetically heterogeneous, and so were streak purified. Individual colonies were screened for retention of markers and the suppressive phenotype of increased spore viability. Genomic DNA was extracted and used to transform the *gerA\** locus into the parental Δ*5* background. All backcrossed mutants retained the suppressive phenotype, indicating linkage to *gerA\**. The *gerA\** locus was subjected to PCR followed by Sanger sequencing to determine mutations.

### Spore purification with lysozyme and SDS

25 mL of overnight DSM culture was collected and washed three times with sterile water. Spores were resuspended in PBS containing lysozyme at a concentration of 1.5 mg/mL. Spores were then incubated at 37°C with agitation for one hour. SDS was added to a final concentration of 2% (w/v) and spores were incubated for an additional 30 minutes at 37°C with agitation. Spores were then washed five times with water to remove the SDS.

### Spores purified with density gradient

25 mL of overnight DSM culture were heat-treated (80 °C for 20 min) and washed three times with sterile water. The spore pellet was resuspended in 1 ml of 20% (w/v) Histodenz (Sigma) and incubated for 30 min on ice. This suspension was then gently pipetted on top of 2 mL of 40% (w/v) Histodenz which itself was layered on 6 ml of 50% (w/v) Histodenz. The gradient was centrifuged at 4,000 RPM for 90 min at 4 °C and the supernatant, which contained phase-dark spores, vegetative cells, and cell debris, was siphoned off. The pellet was washed three times with sterile water. Pellets were suspended in 1 mL H_2_O and kept at 4 °C All spore preparations were evaluated by phase-contrast microscopy and contained >99% phase bright spores.

### Microscopy

Overnight sporulation cultures and purified spores were concentrated by centrifugation and then immobilized on pads made of 2% (w/v) agarose in PBS. Phase-contrast microscopy was performed using a Nikon TE2000 inverted microscope, Nikon Intensilight Metal Halide Illumination, a Photometrics CoolSNAP HQ2 monochrome CCD camera, and a Plan Apo 100x/1.4 oil Ph3 DM objective. All exposure times were 250 ms. Image acquisition was performed using Nikon Elements Acquisition Software AR 3.2. Image analysis and processing were performed in Metamorph.

### SDS-PAGE and Immunoblotting

Spores were purified with lysozyme and SDS and then suspended in 0.4 mL PBS with protease inhibitors. Spore resuspensions were added to 2 mL tubes containing Lysing Matrix B (MP Biomedicals, Irvine, CA), and chilled on ice. Spores were then ruptured mechanically using FastPrep (MP Biomedicals) at 6.5 m/s for 1 minute. 0.4 mL of 2x Laemmli sample buffer with 10% (v/v) β-mercaptoethanol was immediately added and tubes were vortexed. Samples were then incubated at 80 °C for 5 minutes and centrifuged at maximum speed for 10 minutes. Supernatants were collected and total protein was normalized using the Non-Interfering Protein Assay (G-Biosciences, St. Louis, MO).

All samples were separated by SDS-PAGE on 17.5% resolving gels, electroblotted onto Immobilon-P membranes (Millipore, Burlington, MA), and blocked in 5% nonfat milk in PBS with 0.5% Tween-20. Membranes were then probed with anti-GerAA (1:5000) (30), anti-SigA (1:10,000) (31), or anti-SleB (1:5,000) (32)diluted in 3% BSA in PBS with 0.05% Tween-20. Primary antibodies were detected using horseradish-peroxidase conjugated anti-rabbit antibodies (BioRad) and detected with Western Lightning ECL reagents.

### Measuring DPA levels

Spores were purified using lysozyme and SDS and resuspended to OD_600_ = 1 in water. Spore suspensions were then incubated at 100 °C for 30 minutes to release DPA. Suspensions were subjected to centrifugation and supernatants were added to a solution at a final concentration of 100 μM TbCl_3_ in 1M sodium acetate, pH 5.6. Fluorescence was measured at 545 nm with excitation set at 272 nm. Each sample was analyzed in technical triplicate and compared to a standard curve to determine DPA concentration.

### Germination assays

Spores were purified by Histodenz gradient and resuspended in 25 mM HEPES pH 7.4 at OD_600_ = 1.2. Suspensions were heat activated for 30 minutes at 70 °C followed by 20 minutes on ice. Suspensions were then transferred to a clear, 96-well, flat-bottom tray, and either buffer (25 mM HEPES pH 7.4) or 1mM L-alanine in buffer (25 mM HEPES pH 7.4) was added for a final OD_600_ = 0.6. The 96-well trays were agitated in a TECAN plate reader at 37 °C and optical density readings were taken every two minutes. Spores were tested in technical triplicate and values were averaged.

## Acknowledgements

We thank all members of the Bernhardt-Rudner super-group past and present for helpful advice, discussions, and encouragement. We thank Paula Montero Llopis, Ryan Stephansky, and the HMS Microscopy Resources on the North Quad (MicRoN) core at Harvard Medical School for advice on microscopy. Support for this work comes from GM086466, GM127399 and funds from the HMS Dean’s Initiative (DZR). JDA was funded by the National Institutes of Health Grant F32GM130003. LA is a Simons Foundation fellow of the Life Sciences Research Foundation.

