## Supplemental Data for "Genetic evidence for signal transduction within the *Bacillus subtilis* GerA germinant receptor"

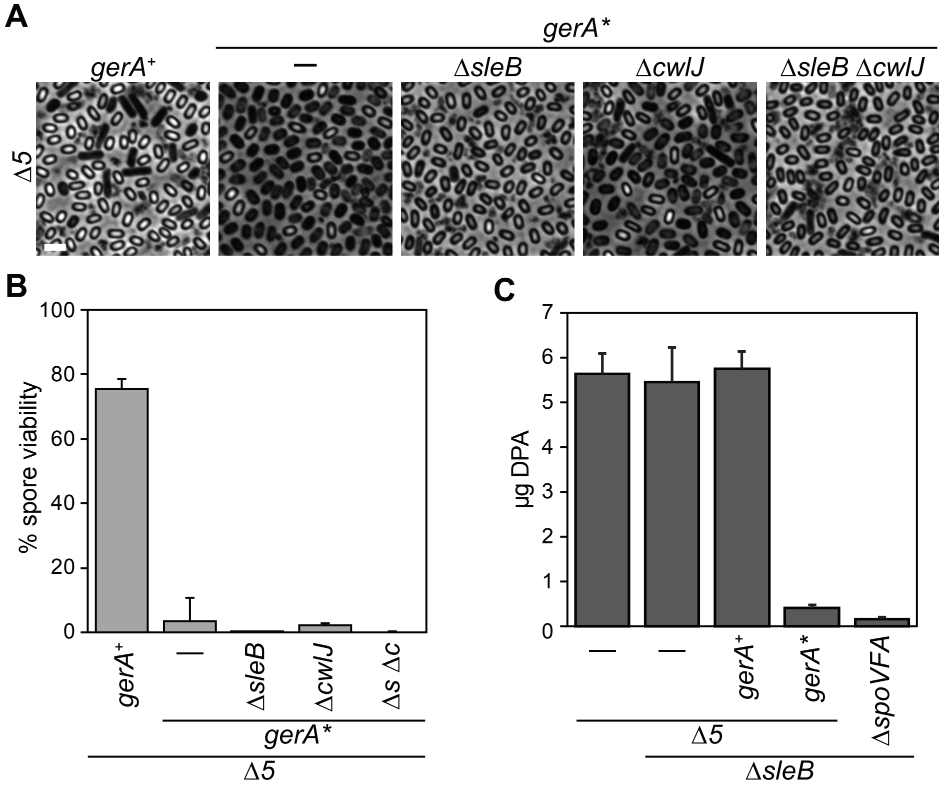


**Supplemental Figure 1: GerA* triggers DPA release and SleB activation when it is the sole germinant receptor.** (A) Phase-contrast micrographs of cultures sporulated by nutrient exhaustion. *Δ5* – *ΔgerA ΔgerBB ΔgerKB ΔyndE ΔyfkT*. Experiments were performed in biological triplicate; representative images are shown. Scale bar is 2μm. (B) Sporulated cultures in (A) were heat treated (80°C for 20 min), and serial dilutions were plated on LB to assess heat-resistant colony forming units. Wild type spore viability (3.3x108 CFU/ml) was set to 100%. *Δs Δc* – *ΔsleB ΔcwlJ*. Error bars indicate standard deviation, n=3. (C) Phase-grey and -bright spores were purified from sporulated cultures in (A) using lysozyme and SDS. Spores were boiled to release DPA. DPA was then quantified by measuring fluorescence when combined with a solution containing $TbCl_3$ compared to standards. Values are reported as micrograms of DPA released from 1ml of purified spores adjusted to $OD_{600}$=1. Error bars indicate standard deviation, n=4.

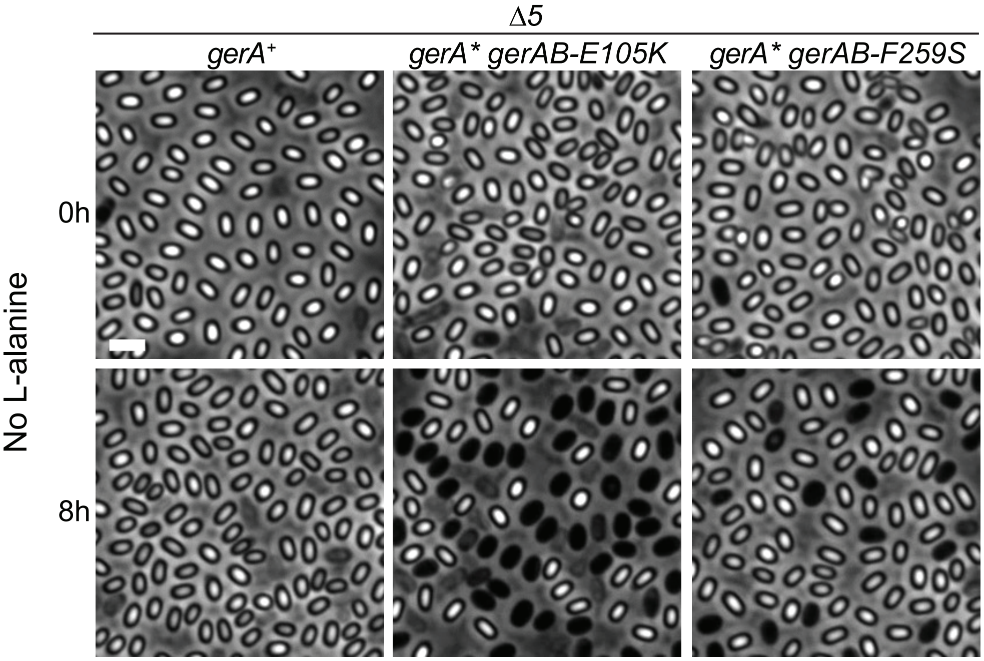


**Supplemental Figure 2: Constitutive germination of spores harboring *gerAA(P326S)* and *gerAB(E105K)* or *gerAB(F259S)*.** Phase-contrast micrographs of purified spores before and after agitation in buffer for 8 h at 37°C. Images correspond to the plots in Figures 3C and 3D. Representative images of three biological triplicates are shown. Scale bar is 2μm.

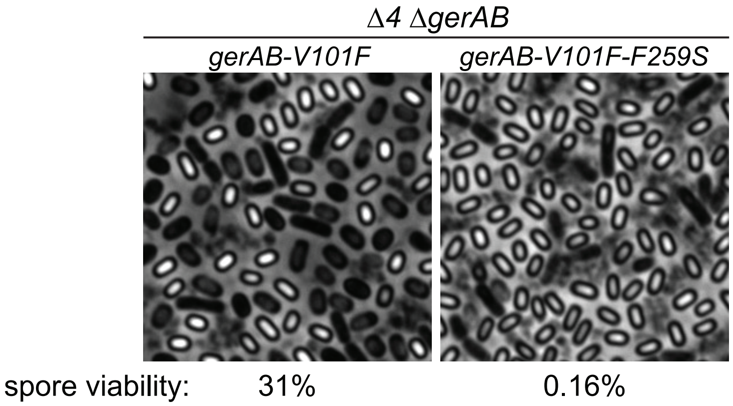


**Supplemental Figure 3: Epistatic analysis of gerAB-F259S and gerAB-V101F.** Cultures were sporulated by nutrient exhaustion. Representative phase-contrast micrographs of three biological replicates are shown. Scale bar is 2μm. Cultures were heat-treated (80°C for 20 min) then serially diluted and plated on LB to assess heat-resistant colony forming units. *gerAB*$^{+}$ spore viability (3.7 x108 CFU/ml) was set to 100%.

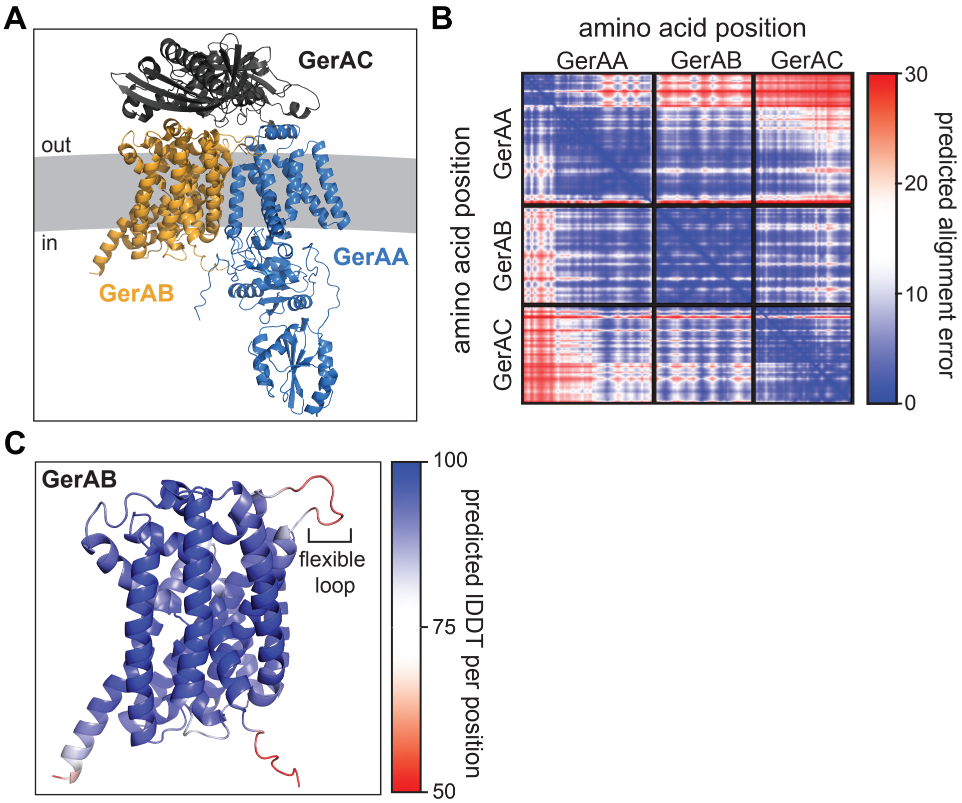


**Supplemental Figure 4: Structural prediction of GerA complex.** (A) Alphafold2-predicted structures of GerAA, GerAB, and GerAC in complex, situated in the inner spore membrane (grey). (B) Predicted alignment error of all residues against all residues. Low error (blue) corresponds to well-defined relative domain positions. (C) Predicted lDDT per position mapped onto the predicted GerAB structure. Higher plDDT (blue) corresponds to a more confident prediction.

**Supplemental Table 1.** Suppressors of *gerAA-P326S*

| GerAA Alteration | n | GerAB Alteration | n | GerAC Alteration | n |
|---|---|---|---|---|---|
| P126L | 1 | E105K | 1 | S28I | 1 |
| M165I | 1 | R107Q | 1 | V119F | 1 |
| A236S | 1 | R107W | 3 | S342P | 2 |
| M263I | 1 | V163E | 1 | T368L | 1 |
| S265P | 1 | W253L | 1 | T368R | 1 |
| L293V | 1 | **F259S** | **1** | | |
| S294P | 1 | **F259L** | **2** | | |
| **A299V*** | **1** | **G266D** | **2** | | |
| **S302T*** | **1** | **G266S** | **6** | | |
| **A313T** | **3** | **I267R** | **1** | | |
| **A313V** | **2** | | | | |
| **T315M** | **2** | | | | |
| **S317L** | **1** | | | | |
| **A318T** | **1** | | | | |
| **A318V** | **3** | | | | |
| **E321G** | **4** | | | | |
| **E321A** | **1** | | | | |
| **E321V** | **1** | | | | |
| **P324L** | **1** | | | | |
| **S326P** | **6** | | | | |
| T337I | 1 | | | | |
| Q366H | 1 | | | | |
| V369I | 1 | | | | |
| E370A | 1 | | | | |
| A371V | 1 | | | | |
| L378F | 3 | | | | |
| A386T | 8 | | | | |
| T391I | 1 | | | | |
| F401V | 1 | | | | |
| R402Q | 3 | | | | |
| S408P | 3 | | | | |
| **Total** | 58 | | 19 | | 6 |

**Bold** indicates alteration is within the extracellular loops discussed in the manuscript

*Residues 299 and 302 of GerAA are variable across laboratory strains. See reference 25 for further details

**Supplemental Table 2.** *Bacillus subtilis* strains used in this study

| Strain | Genotype | Source | Figure |
|---|---|---|---|
| 168 | *trpC2* | Zeigler *et al.*, 2008 | 1, 2, 3, S1 |
| BJA153a | *ycgO::gerAA-gerAB-gerAC (spec)* | This study | 1 |
| BJA134b | *ycgO::gerAA(P326S)-gerAB-gerAC (spec)* | This study | 1 |
| BJA567 | *ycgO::gerAA(P326S)-gerAB-gerAC (spec)* Δ*sleB::lox72* | This study | 1 |
| BJA148a | *ycgO::gerAA(P326S)-gerAB-gerAC (spec)* Δ*cwlJ::lox72* | This study | 1 |
| BJA568 | *ycgO::gerAA(P326S)-gerAB-gerAC (spec)* Δ*sleB::lox72* Δ*cwlJ::lox72* | This study | 1 |
| BDR3487 | Δ*sleB::lox72* | Amon *et al.,* 2020 | 1, S1 |
| BDR4199 | Δ*sleB::lox72* Δ*spoVFA:lox72* | This study | 1, S1 |
| BLA176 | Δ*gerA::cat* Δ*gerBB::lox72* Δ*gerKB::lox72* Δ*yfkT::lox72* Δ*yndE::lox72 ycgO::kan* | Artzi, *et al.* 2021 | 2 |
| BLA197 | Δ*gerA::cat* Δ*gerBB::lox72* Δ*gerKB::lox72* Δ*yfkT::lox72* Δ*yndE::lox72 ycgO::gerAA-gerAB-gerAC (spec)* | Artzi, *et al.* 2021 | 2, 3, S1, S2 |
| BJA177a | Δ*gerA::cat* Δ*gerBB::lox72* Δ*gerKB::lox72* Δ*yfkT::lox72* Δ*yndE::lox72 ycgO::gerAA(P326S)-gerAB-gerAC (spec)* | This study | 2, S1 |
| BJA185f | Δ*gerA::cat* Δ*gerBB::lox72* Δ*gerKB::lox72* Δ*yfkT::lox72* Δ*yndE::lox72 ycgO::gerAA(A313T, P326S)-gerAB-gerAC (spec)* | This study | 2 |
| BJA185k | Δ*gerA::cat* Δ*gerBB::lox72* Δ*gerKB::lox72* Δ*yfkT::lox72* Δ*yndE::lox72 ycgO::gerAA(P326S, A386T)-gerAB-gerAC (spec)* | This study | 2 |
| BJA561 | Δ*gerA::cat* Δ*gerBB::lox72* Δ*gerKB::lox72* Δ*yfkT::lox72* Δ*yndE::lox72 ycgO::gerAA(P326S)-gerAB(E105K)-gerAC (spec)* | This study | 2, 3, S2 |
| BJA185d | Δ*gerA::cat* Δ*gerBB::lox72* Δ*gerKB::lox72* Δ*yfkT::lox72* Δ*yndE::lox72 ycgO::gerAA(P326S)-gerAB(F259S)-gerAC (spec)* | This study | 2, 3, S2 |
| BJA235 | Δ*gerA::cat* Δ*gerBB::lox72* Δ*gerKB::lox72* Δ*yfkT::lox72* Δ*yndE::lox72 ycgO::gerAA(P326S)-gerAB-gerAC(S28I) (spec)* | This study | 2 |
| BJA236 | Δ*gerA::cat* Δ*gerBB::lox72* Δ*gerKB::lox72* Δ*yfkT::lox72* Δ*yndE::lox72 ycgO::gerAA(P326S)-gerAB-gerAC(S342P) (spec)* | This study | 2 |
| BLA174 | Δ*gerAB::lox72* Δ*gerBB::lox72* Δ*gerKB::lox72* Δ*yfkT::lox72* Δ*yndE::lox72 ycgO::kan* | Artzi, *et al.* 2021 | 3 |
| BLA178 | Δ*gerAB::lox72* Δ*gerBB::lox72* Δ*gerKB::lox72* Δ*yfkT::lox72* Δ*yndE::lox72 ycgO::gerAB* | Artzi, *et al.* 2021 | 3, 4 |
| BJA278 | Δ*gerAB::lox72* Δ*gerBB::lox72* Δ*gerKB::lox72* Δ*yfkT::lox72* Δ*yndE::lox72 ycgO::gerAB(E105K)* | This study | 3 |
| BJA279 | Δ*gerAB::lox72* Δ*gerBB::lox72* Δ*gerKB::lox72* Δ*yfkT::lox72* Δ*yndE::lox72 ycgO::gerAB(R107Q)* | This study | 3 |
| BJA280 | Δ*gerAB::lox72* Δ*gerBB::lox72* Δ*gerKB::lox72* Δ*yfkT::lox72* Δ*yndE::lox72 ycgO::gerAB(R107W)* | This study | 3 |
| BJA281 | Δ*gerAB::lox72* Δ*gerBB::lox72* Δ*gerKB::lox72* Δ*yfkT::lox72* Δ*yndE::lox72 ycgO::gerAB(W253L)* | This study | 3 |
| BJA282 | Δ*gerAB::lox72* Δ*gerBB::lox72* Δ*gerKB::lox72* Δ*yfkT::lox72* Δ*yndE::lox72 ycgO::gerAB(F259S)* | This study | 3, 4 |
| BJA283 | Δ*gerAB::lox72* Δ*gerBB::lox72* Δ*gerKB::lox72* Δ*yfkT::lox72* Δ*yndE::lox72 ycgO::gerAB(G266D)* | This study | 3 |
| BJA284 | Δ*gerAB::lox72* Δ*gerBB::lox72* Δ*gerKB::lox72* Δ*yfkT::lox72* Δ*yndE::lox72 ycgO::gerAB(G266S)* | This study | 3 |
| BJA285 | Δ*gerAB::lox72* Δ*gerBB::lox72* Δ*gerKB::lox72* Δ*yfkT::lox72* Δ*yndE::lox72 ycgO::gerAB(I267R)* | This study | 3 |
| BJA569 | Δ*gerAB::lox72* Δ*gerBB::lox72* Δ*gerKB::lox72* Δ*yfkT::lox72* Δ*yndE::lox72 ycgO::gerAB(V101F)* | This study | S3 |
| BJA570 | Δ*gerAB::lox72* Δ*gerBB::lox72* Δ*gerKB::lox72* Δ*yfkT::lox72* Δ*yndE::lox72 ycgO::gerAB(T287L)* | This study | 4 |
| BJA571 | Δ*gerAB::lox72* Δ*gerBB::lox72* Δ*gerKB::lox72* Δ*yfkT::lox72* Δ*yndE::lox72 ycgO::gerAB(V101F, F259S)* | This study | S3 |
| BJA572 | Δ*gerAB::lox72* Δ*gerBB::lox72* Δ*gerKB::lox72* Δ*yfkT::lox72* Δ*yndE::lox72 ycgO::gerAB(F259S,T287L)* | This study | 4 |

| | | | |
|---|---|---|---|
| BJA317a | *ΔgerA::cat ΔgerBB::lox72 ΔgerKB::lox72 ΔyfkT::lox72 ΔyndE::lox72 ycgO::gerAA(P326S)-gerAB-gerAC (spec) ΔsleB::erm* | This study | S1 |
| BJA318a | *ΔgerA::cat ΔgerBB::lox72 ΔgerKB::lox72 ΔyfkT::lox72 ΔyndE::lox72 ycgO::gerAA(P326S)-gerAB-gerAC (spec) ΔcwlJ::erm* | This study | S1 |
| BJA319a | *ΔgerA::cat ΔgerBB::lox72 ΔgerKB::lox72 ΔyfkT::lox72 ΔyndE::lox72 ycgO::gerAA-gerAB-gerAC (spec) ΔsleB::erm* | This study | S1 |
| BJA568 | *ΔgerA::cat ΔgerBB::lox72 ΔgerKB::lox72 ΔyfkT::lox72 ΔyndE::lox72 ycgO::gerAA(P326S)-gerAB-gerAC (spec) ΔsleB::lox72 ΔcwlJ::lox72* | This study | S1 |
| BJA287a | *ΔgerA::cat ΔgerBB::lox72 ΔgerKB::lox72 ΔyfkT::lox72 ΔyndE::lox72 ycgO::kan ΔsleB::erm* | This study | S1 |

**Supplemental Table 3.** Plasmids used in this study

| Plasmid | Description | Source |
|---|---|---|
| pLA25 | *ycgO::gerAA-gerAB-gerAC (spec)* | Artzi, *et al*. 2021 |
| pJA39 | *ycgO::gerAA(P326S)-gerAB-gerAC (spec)* | This study |
| pLA13 | *ycgO::gerAB (spec)* | Artzi, *et al*. 2021 |
| pJA048 | *ycgO::gerAB(E105K) (spec)* | This study |
| pJA049 | *ycgO::gerAB(R107Q) (spec)* | This study |
| pJA050 | *ycgO::gerAB(R107W) (spec)* | This study |
| pJA051 | *ycgO::gerAB(W253L) (spec)* | This study |
| pJA052 | *ycgO::gerAB(F259S) (spec)* | This study |
| pJA053 | *ycgO::gerAB(G266D) (spec)* | This study |
| pJA054 | *ycgO::gerAB(G266S) (spec)* | This study |
| pJA055 | *ycgO::gerAB(I267R) (spec)* | This study |
| pLA129 | *ycgO::gerAB(V101F) (spec)* | Artzi, et al. 2021 |
| pJA078 | *ycgO::gerAB(V101F, F259S) (spec)* | This study |
| pLA125 | *ycgO::gerAB(T287L) (spec)* | Artzi, et al. 2021 |
| pJA079 | *ycgO::gerAB(F259S, T287L) (spec)* | This study |

**Supplemental Table 4.** Oligonucleotides used in this study

| Oligonucleotide | Sequence |
|---|---|
| oLA111 | ggcACTAGTatcctttgaatatttgtattttggatttg |
| oLA108 | ggcGGATCCctggatgtcagtgaccggacg |
| oJA105 | ggcgaacagggaaaacgtgccgttctctccgatatttgaagccctgctgatg |
| oJA106 | catcagcagggcttcaaatatcggagagaacggcacgttttccctgttcgcc |
| oJA152 | TCCTCGGCGTAGCCAGCTTCAAGACACGGGCAATGGCTGA |
| oJA153 | TCAGCCATTGCCCGTGTCTTGAAGCTGGCTACGCCGAGGA |
| oJA154 | GCGTAGCCAGCTTCGAGACATGGGCAATGGCTGAAATGGTGA |
| oJA155 | TCACCATTTCAGCCATTGCCCATGTCTCGAAGCTGGCTACGC |
| oJA156 | CGTAGCCAGCTTCGAGACACAGGCAATGGCTGAAATGGTGA |
| oJA157 | TCACCATTTCAGCCATTGCCTGTGTCTCGAAGCTGGCTACG |
| oJA158 | CGAGGTGAAAACGCTGATTTTGCCGACTATTTCTCTCTTTC |
| oJA159 | GAAAGAGAGAAATAGTCGGCAAAATCAGCGTTTTCACCTCG |
| oJA160 | TTGGCCGACTATTTCTCTCTCTCAGTCCTTTGAGCTTAAAG |
| oJA161 | CTTTAAGCTCAAAGGACTGAGAGAGAGAAATAGTCGGCCAA |
| oJA162 | TCAGTCCTTTGAGCTTAAAGACATATTTATTGAACGGTTTG |
| oJA163 | CAAACCGTTCAATAAATATGTCTTTAAGCTCAAAGGACTGA |
| oJA164 | TTCAGTCCTTTGAGCTTAAAAGCATATTTATTGAACGGTTT |
| oJA165 | AAACCGTTCAATAAATATGCTTTTAAGCTCAAAGGACTGAA |
| oJA166 | GTCCTTTGAGCTTAAAGGCAGATTTATTGAACGGTTTGAAT |
| oJA167 | ATTCAAACCGTTCAATAAATCTGCCTTTAAGCTCAAAGGAC |
